# Improving localization precision via restricting biomolecule confined stochastic motion in SMLM

**DOI:** 10.1101/2021.03.16.435087

**Authors:** Jielei Ni, Bo Cao, Gang Niu, Tingying Xia, Danni Chen, Wanlong Zhang, Yilin Zhang, Xiaocong Yuan, Yanxiang Ni

**Affiliations:** Nanophotonics Research Center, Shenzhen Key Laboratory of Micro-Scale Optical Information Technology & Institute of Microscale Optoelectronics, College of Physics and Optoelectronic Engineering, College of Electronics and Information Engineering, Shenzhen University, Shenzhen 518060, China; Phil Rivers Technology, Beijing 100871, China

**Keywords:** localization precision, single molecule localization microscopy, stochastic optical reconstruction microscopy

## Abstract

Single-molecule localization microscopy (SMLM) boosts its applications when combined with the studies of cells, in which nanometer-sized biomolecules are irresolvable due to diffraction limit unless being subjected to SMLM. Although being well immobilized, given the nanometer sizes of biological molecules, they are still capable of movement stochastically around their immobilized sites. The influence of such motion on image quality and possible improvements have not yet been systematically investigated. Here, we accessed the biomolecule stochastic motion in SMLM by calculating the displacements between different localizations from the same molecule in single-molecule samples of Alexa Fluor-647-conjugated oligonucleotides. We found that, for most molecules, localization displacements at random frame intervals are remarkably larger than those between temporally neighbouring frames despite of drift correction, showing that biomolecule stochastic motion is involved in SMLM. Furthermore, the localization displacements were observed to increase with frame intervals and then saturate, suggesting biomolecule stochastic motion is confined within a finite area. Moreover, we showed that the localization precision is deteriorated by enlarging molecule sizes and improved by sample post-fixation. This study reveals confined stochastic motion of biomolecules increase localization uncertainty in SMLM, and improved localization precision can be achieved via restricting biomolecule stochastic motion.

## 1 Introduction

Single-molecule localization microscopy (SMLM), such as stochastic optical reconstruction microscopy (STORM) [1] and photoactivation localization microscopy (PALM) [2], has circumvented the diffraction barrier and offered lateral resolution of 15-25 nm [1–4]. Due to the unmatched resolution among all fluorescence microscopies and its non-invasive property to biological samples, SMLM plays an irreplaceably essential role in biological studies, one important aim for which is to visualize nanoscale-sized biomolecules including proteins, DNA, and lipids in cells [5–6].

SMLM provides super-resolution via stochastically activating single photo-switchable fluorophores and accurately determining their localizations. To visualize the intracellular ultra-structures, biomolecules of interest are commonly labelled with switchable fluorophores via covalent conjugation [7–8] or genetically encoding protein fusion [9–10]. To maintain subcellular structures, biological samples are generally immobilized at the beginning of sample preparation, so that most biomolecules cannot diffuse but retain at original positions except some transmembrane and membrane-bound proteins that require special fixation [11–12]. The precision of localizing single molecules is mainly dominated by the number of photons collected from the activating fluorophore, such as Alexa Fluor-647 (AF647) [13–15]. However, in light of the high resolution and long imaging time, any motion during acquisition probably enhances the localization uncertainty and hampers image resolution. In this regard, intense studies have been focused on sample drift correction to ensure the high quality of final super-resolution image [16–18]. Since biomolecules as well as the antibodies or probes binding to them exhibit sizes of one to a few tens of nanometers [7–8, 19], confined stochastic motion of immobilized biomolecules is expected to exist in SMLM as the imaging is performed in buffer and commonly at room temperature. Given the high spatial resolution and the relatively long imaging time of SMLM, such a stochastic motion during the acquisition probably decreases localization precision. However, it has not been evident with experiment data whether confined stochastic motion is detectable in SMLM or affects localization precision

Studies of confined biomolecule stochastic motion has been limited for several reasons. Firstly, the stochastic activation of fluorophores offers discontinuous trajectory that is far insufficient for tracking the biomolecule motion. Secondly, due to the sample immobilizations, motion is confined within a finite area, the range of which is comparable to the spatial resolution of SMLM. This makes the motion challenging to be resolved. Thirdly, due to dense labeling in regular biological samples, it is almost impossible to identify motion of a target biomolecule or the specific probe or antibody binding to it. Therefore, new analysis is required for clarifying the detectability and impact of immobilized biomolecules in SMLM, so that effective strategies might be applied to improve image resolution.

In this paper, to address the questions about confined biomolecule stochastic motion, we applied single-molecule samples containing sparsely-immobilized AF647-conjugated oligonucleotides. The sizes of these oligomers are in the range of regular biomolecule sizes. We assessed the stochastic motion of the immobilized molecules based on STORM imaging data. Briefly, we calculated the displacements between two localizations at different frame intervals from the same molecule. Z_1_-score for each molecule was proposed to statistically evaluate the significance of localization displacements between temporally neighboring frames compared to those between two random frames. It turned out that, for most of molecules examined, the z_1_-score is negative despite of sample drift correction, showing the stochastic motion of immobilized molecules during sequential imaging. Furthermore, in regard to different frame intervals, the localization displacements were observed to increase with intervals and then saturate, suggesting molecule motion is confined within a finite area. Moreover, localization precision was found to be inversely proportional to the amplitude of molecule stochastic motion. We also revealed that enlarging molecule sizes increase localization uncertainty while post-fixation labelled samples improved localization precision, which implies strategy for optimizing localization precision via minimizing stochastic motion of immobilized biomolecules in regular biological samples.

## 2 Results and discussion

### 2.1 Confined stochastic motion in modal sample

To assess the potential stochastic motion of immobilized biomolecules in STORM, we apply in this study a single molecule sample containing 40-nucleotide oligomers, which are of similar sizes to and used here to mimic target molecules as well as their antibodies/probes in biological samples. Each oligonucleotide is conjugated to an AF647 molecule at 5’ end for reporting its position and immobilized on gelatin-coated glass via nonspecific binding of biotin at 3’ end to gelatin (**Figure 1a**). In light of their nanometer sizes, we hypothesize that AF647-conjugated oligonucleotides in imaging buffer stochastically move around the immobilized sites within a confined region under the control of a combined strain from the immobilized site and the thermal motion of surrounding liquid molecules. Molecule motion (the red dashed circle) is recorded in a series of frames (Fn, Fn+1, Fn+2, …, Fn+x) among an image sequence in STORM (**Figures 1b1 an 1b2**), although the information is insufficient for outlining its trajectory. In regard to sample drift occurring between two frames (Fn and Fn+x), fluorescent bead can be used to correct the position back to the original Fn (**Figure 1c**, green dashed arrow). Meanwhile, molecule stochastic motion (blue arrow) is not corrected by drift compensation, resulting in a new position (marked as Fn+x) instead of the original position at Fn. Therefore, the series of positions/localizations after drift correction can provides essential clues for assessing stochastic motion.

**Figure 1.**
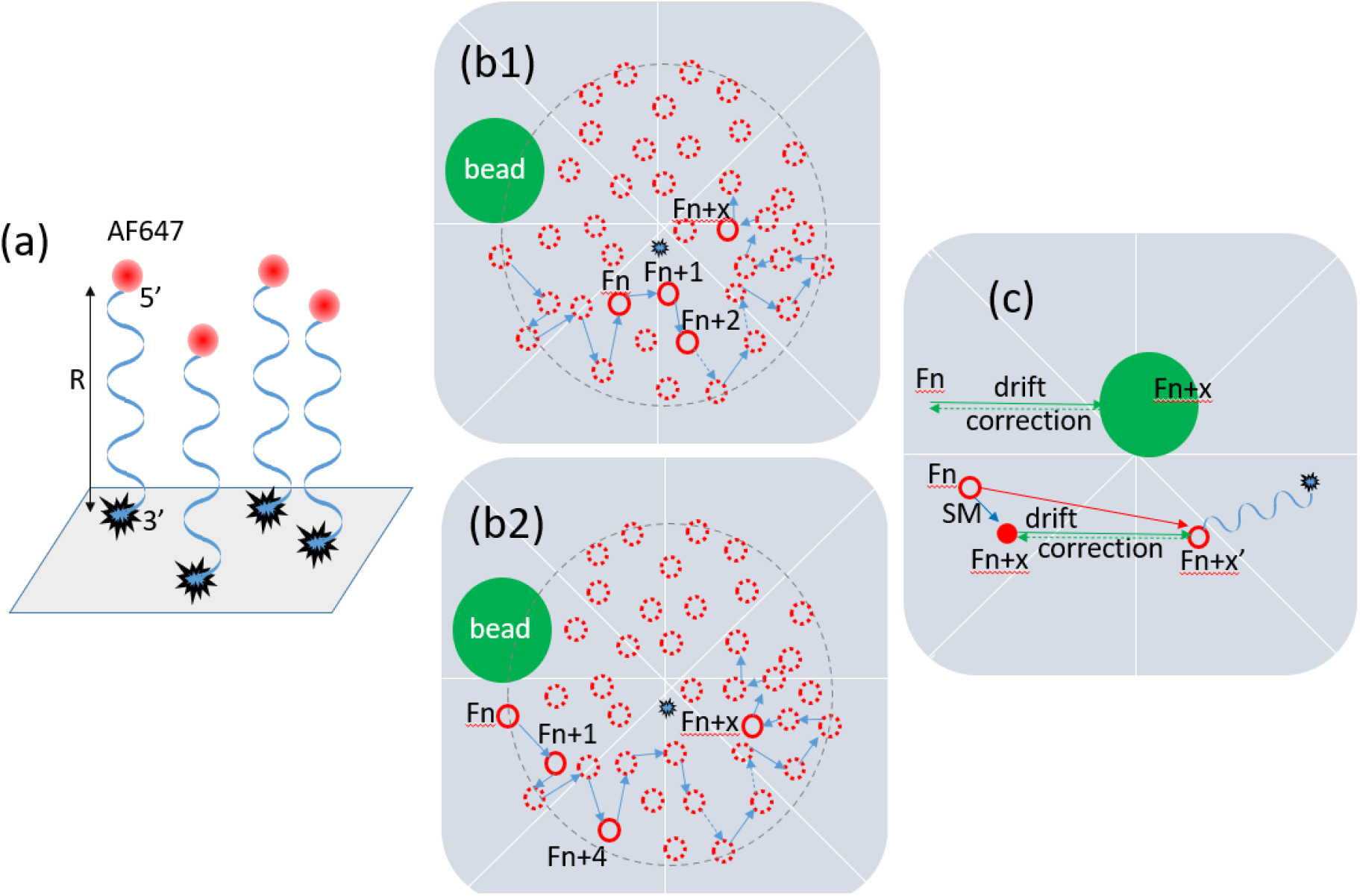
Illustration of confined stochastic motion of immobilized oligonucleotides mimicking biomolecules in STORM. (a) The 40 nucleotide oligomers, each of which is conjugated to a single AF647 (red dot) at 5’ end and to a biotin at 3’ end, are sparsely immobilized on gelatin-coated glass surface via biotin non-specific binding. The oligonucleotide molecule is expected to exhibit a size of around 12 nm (R) to mimic target biomolecules as well as their oligomer probes or antibodies in regular STORM samples. During acquisition, the molecules in imaging buffer are supposed to undergo stochastic motion within a constrained cycle region (gray dashed line), which takes the immobilized site as center and R as radius. (b1 and b2) Due to the stochastic activation of switchable fluorophore, the immobilized oligonucleotide molecule undergoing confined motion is randomly recorded in different frames (Fn, Fn+1, …, Fn+x) during sequential imaging. Although the AF647 molecule can move to any position inside the confined cycle region, some molecules might mainly move around the central region (b1) while others might cover a wider area (b2). Filled green cycle represents fluorescent bead for sample drift correction. (c) Position change (red arrow) of a certain molecule between Fn and Fn+x is likely a combination of its confined stochastic motion (blue arrow) and sample drift (green arrow). The part of change due to sample drift can be corrected via comparing the position alteration of fluorescent bead from Fn to Fn+x, while confined stochastic motion cannot be corrected by drift correction. AF647, Alexa Fluor-647; Fn, Frame n; Fn+x, frame n+x; SM, stochastic motion.

### 2.2 Using Z-score to study confined stochastic motion

To address if stochastic motion of immobilized biomolecules is existing and detected in SMLM, we collected STORM imaging data from the single molecule samples and calculated for each single molecule the displacements between any two of its localizations. Then, *z*_1_-score was used to assess the significance of the localization displacements between two temporally neighboring frames compared to those between two random frames for each single molecule. The calculation of *z*_1_-score is illustrated in **Figure 2a**.

**Figure 2.**
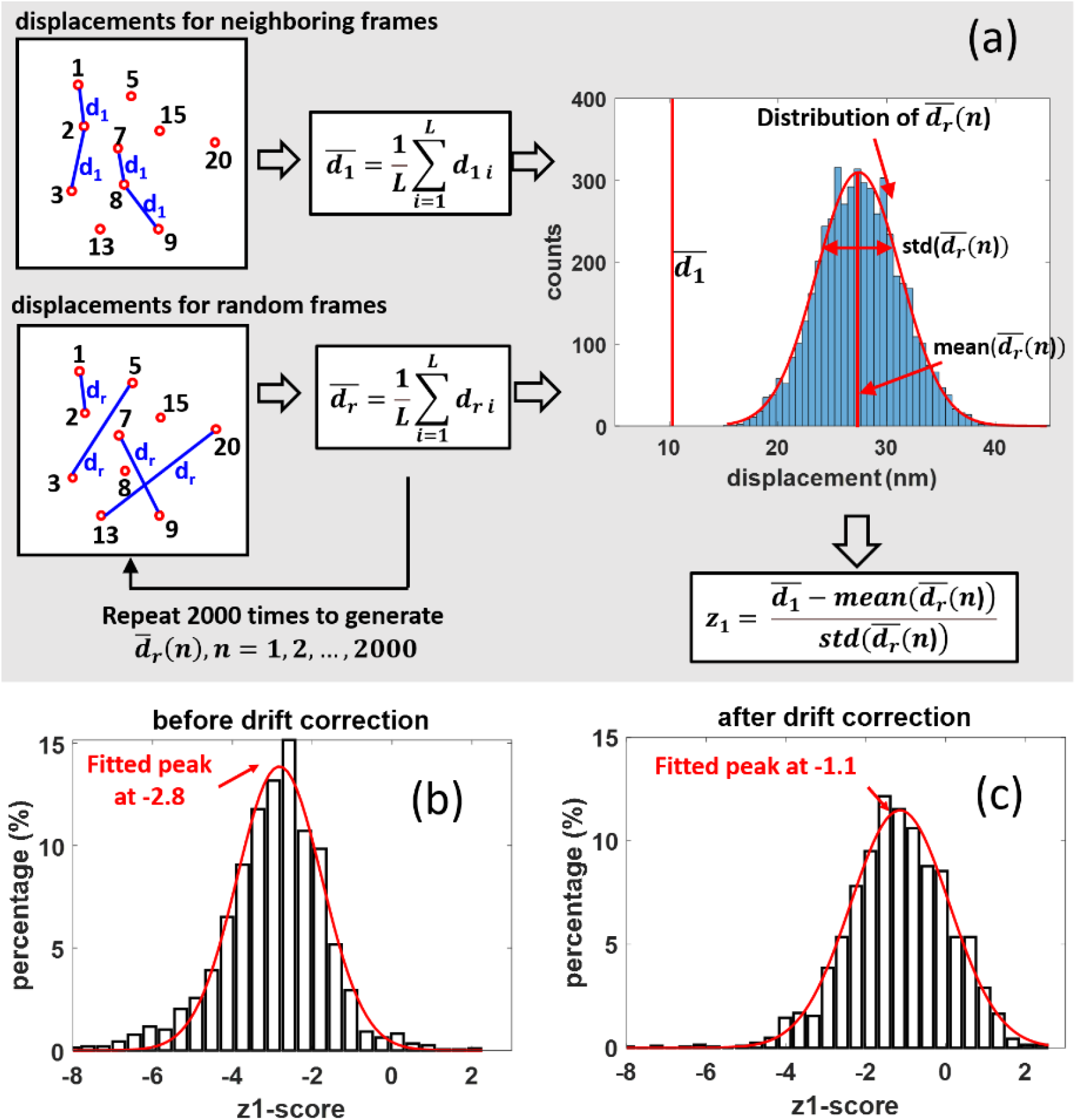
Calculation of neighbouring z-scores of immobilized biomolecule shows the existence of biomolecule stochastic motion after drift correction. (a) Illustration of neighboring z-score (*z*_1_-score) algorithm for a certain molecule. Left-top inset illustrates the displacements *d*_1_ for temporally neighboring localizations. Localizations of a molecule are represented by red circles with frame numbers, from which contain *L* pairs of neighboring frames (*L* = 4 in this example). The displacements *d*_1_ are calculated from the *L* pairs and depicted as blue lines; Left-bottom inset shows one random sampling of localizations with the same number of pairs (sample size, *L*) to calculate *d_r_* (depicted as blue line). The random sampling is repeated 2000 times, each time generating one mean value of *d_r_*. The histogram of the 2000 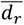 is plotted in the right inset, which shows a normal distribution. The mean value of d, is also plotted as the red line for comparison. The *z*_1_-score determines whether the mean of two sampling groups are statistically significant to each other. (b and c) The distribution of *z*_1_-scores of 2073 molecules, showing a peak at a low value −2.8 before drift correction (b) or at −1.1 after correction (c), respectively.

For each single fluorophore, we defined its localizations as *Loc_i_*(*x_i_,y_i_, f_i_*), *i* = 1, 2, 3,…, *N*, where *N* is the total number of localizations in the 8000 frame-sequence and *f_i_* is the frame number. The displacement for two localizations in temporally neighboring frames in the dataset, which is denoted as *d*_1_

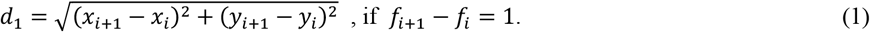

The mean value of all *d*_1_ in the entire dataset can be calculated as

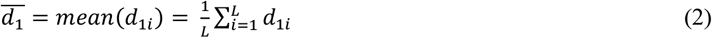

Similarly, the displacement between two localizations in random frames of an image sequence, which is denoted as *d_r_*, can be calculated as,

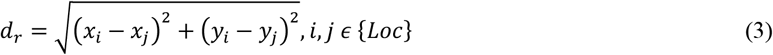

To statistically compare *d*_1_ and *d*_r_, for each molecule we took 2000 random samples from its data set {*Loc*} with the same sample size (*L*) as that in the calculation of *d*_1_, and calculate the mean *d_r_* for each random sample as (**Figure 2a**, left),

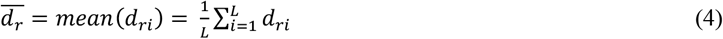

According to Central-Limit Theorem, the distribution of the 2000 sample means would be a normal distribution, as illustrated in the inset (right top) of **Figure 2a**.

To evaluate the statistical difference between *d*_1_ and *d_r_* for a certain molecule, we defined the *z*_1_-score for localizations from temporally neighboring frames as “neighboring z-score”, denoted as *z*_1_ by

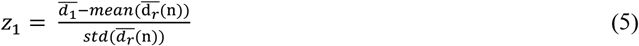

The *z*_1_-score defines how many standard deviations between the neighboring localization displacements and the random localization displacements for a certain molecule. A negative *z*_1_-score reflects that the 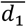 is smaller than the mean of 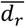, implying molecule motion during acquisition. To assess the potential stochastic movement of immobilized biomolecules during image acquisition, we calculated the *z*_1_-scores both before and after drift correction for 2073 oligonucleotide molecules examined by our STORM imaging. It was found that the *z*_1_-scores of most molecules situate near the peak of −2.8 before correction (**Figure 2b**) and remarkably increase by 1.7 in correction (**Figure 2c**), suggesting effective drift correction. However, the main molecule fraction at a negative *z*_1_-score of around −1.1 after correction (**Figure 2c**) indicates that the molecules are still affected by some other motion, like stochastic motion, which cannot be corrected by the fluorescent bead.

To understand the molecule motion after drift correction, we further analyzed the dynamics of the different z-scores of each molecule at different frame intervals. The displacement for two localizations at an interval of *m* frames in the dataset, which is denoted as *d*_m_, is calculated by revising **Eq. (1)** to

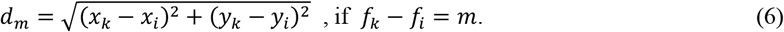

The corresponding z-score, which we defined as “dynamic z-score” and denoted as *z_m_*, is calculated as

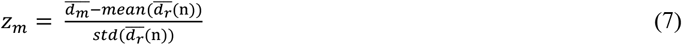

**Figure 3a** shows the ensemble average of *z_m_*-scores over 2073 molecules from our STORM experiment with frame interval *m* increasing from 1 to 30. The dynamic curve starts from the ensemble average of *z*_1_ with a value of −1.1, then increases with frame interval, and finally saturates at around zero after a certain interval. We speculated such behavior to origin from molecule stochastic motion within an area whose boundary is determined by the length of the DNA. At small frame interval, the ensemble averaged displacement increases with time. After a certain time interval when the movement reaches to the entire area, the displacement is restricted by the boundary.

**Figure 3.**
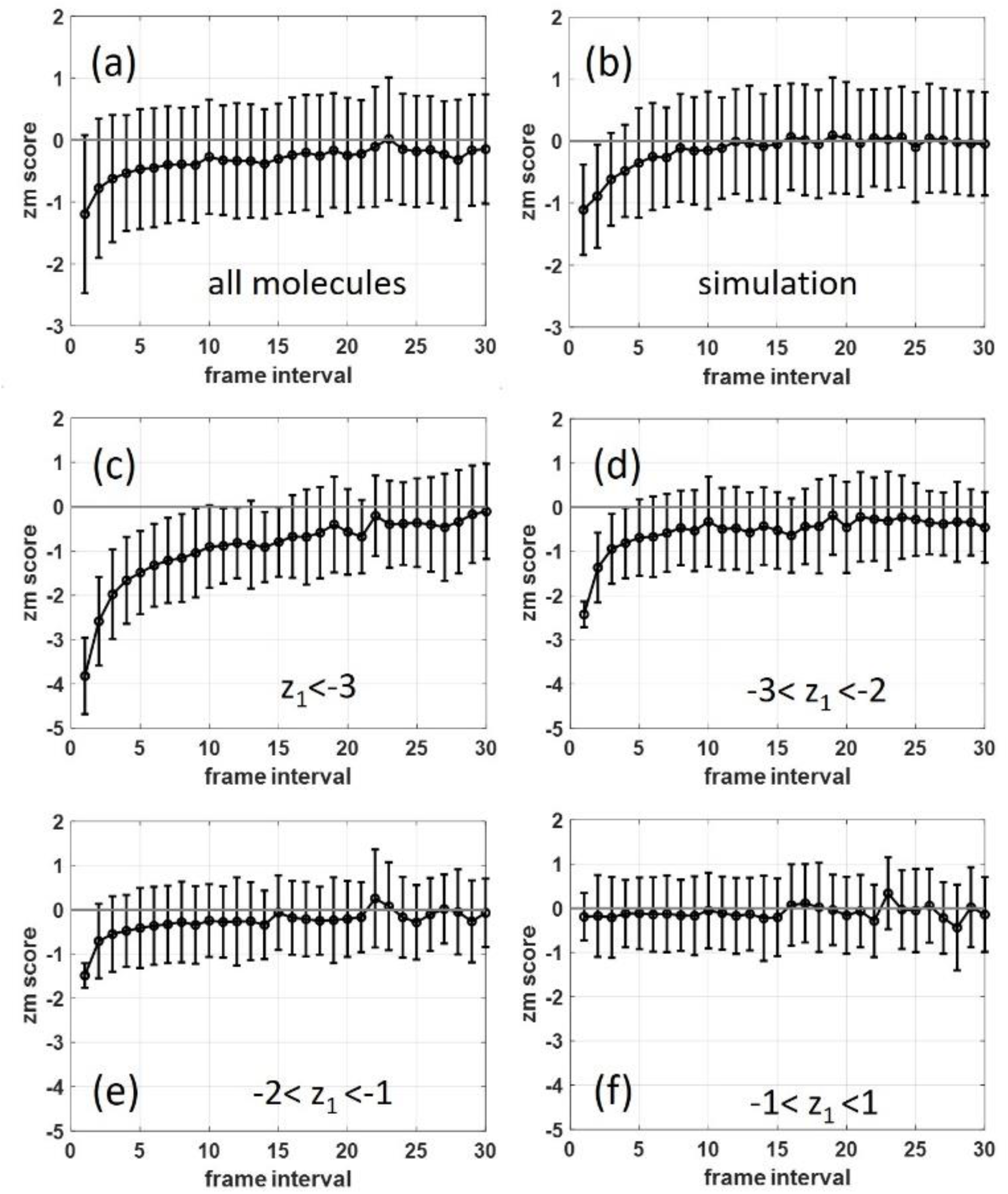
Experimental and theoretical calculations of the dynamic *z_m_*-score after drift correction show the stochastic motion of immobilized bio-molecules within confined regions. (a) The *z_m_*-scores were plotted based on STORM imaging data of 2073 single molecules when frame interval *m* increasing from 1 to 30. Error bar represents the standard deviation of *z_m_*scores of 2073 molecules for a certain interval number *m*. (b) The simulated *z_m_*-scores averaged by 200 molecules show similar saturation pattern when frame interval *m* increasing from 1 to 30. (c-f) The total 2073 molecules are divided into four groups according to different scales of *z*_1_-scores, and their corresponding *z_m_*-scores are separately plotted. For the molecules with a *z*_1_ <-1 (c-e), the *z_m_*-scores increase and saturate at around 0 after *m* reaches a certain number. For the molecules with a *z*_1_-scores between −1 and 1 (f), the *z_m_*-scores randomly fluctuate around zero.

To further confirm our speculation, we simulated the random walk of 200 molecules within a confined circular region with radius of around 13.6 nm (40 *nt* × 0.34 *nm/nt*). In light of their nanometer sizes, biomolecules are expected to be subject to random impulses due to frequent collisions by surrounding liquid molecules and to move continuously but irregularly. Meanwhile, since the biomolecules are immobilized, there is a force restricting the movement within confined area. The details of the simulation are described in **Materials and methods**. The simulation result was depicted in **Figure 3b**. The ensemble average of *z_m_*-scores starts at −1.1, which is in good agreement with the *z*_1_ value of the experimental curve in **Figure 3a**. With increasing frame interval, the *z_m_*-score gradually increases and finally reaches zero. In the experimental curve in Figure 3a, the *z_m_*-score finally saturates to a value below zero, which deviates from the simulation curve and probably can be attributed to residue drift in the experiment. Other possible movements such as vibration was discussed in **Supplementary Materials**.

Although the molecules are subjected to the same imaging buffer and with the same molecule size, due to the random feature of forces by collisions, for different molecules, the motion over a frame interval varies in magnitude and direction. Besides, due to sparse activation, the molecule fluoresces only for a few frames. At different frames, the positions it reported would show different degree of aggregation. As a result, the values of *d_r_* would vary for different molecules. A low *z*_1_-score suggests the molecule having either small *d*_1_ or large *d_r_*. In **Figures 3c-3f,** 2073 molecules from our experiment were grouped according to their *z*_1_-score, and the dynamics of *z_m_*-score of each group were demonstrated. We can see that molecules with lower *z*_1_-score would reach to saturation with a longer frame interval. In **Figure 3f**, the *z_m_*-scores show no dependence on frame interval, indicating the difference between d, and *d_r_* cannot be detected by our system.

### 2.3 Confined stochastic motion decreases localization precision

Due to the high resolution and long acquisition time of STORM, we suspect that molecule stochastic motion would lead to higher localization uncertainty. To analyse the relation between molecule motion and localization precision, molecules from our STORM experiments are grouped according to different *z*_1_-scores for calculating the localization precision. In this calculation, molecules were firstly grouped by their *z*_1_-scores: Group I with 146 molecules with *z*_1_ < −3, Group II with 364 molecules with –3 < *z*_1_, < −2, Group III with 651 molecules with –2 < *z*_1_ < –1, Group IV with 839 molecules with –1 < *z*_1_ < 1, Group V with 68 molecules with 1 < *z*_1_ < 2. To ensure the localization precision for each group were calculated from the same number of molecules, random sampling was performed in this calculation. We take 2000 samples for each group, all of the same size of 30 molecules, and compute the localization precision of each sample. The localization precision was calculated as the standard deviation of localizations. Figure 4a and 4b show the boxplot of localization precision distribution in x-direction and y-direction of each group, respectively. As shown in **Figure 4a**, x-direction localization precision using molecules of different *z*_1_-score groups were inversely correlative to *z*_1_-scores. Similar correlation was observed between y-direction localization precision and *z*_1_-scores (**Figure 4b**). These findings showed that stochastic motion of immobilized biomolecules decreases localization precision.

**Figure 4.**
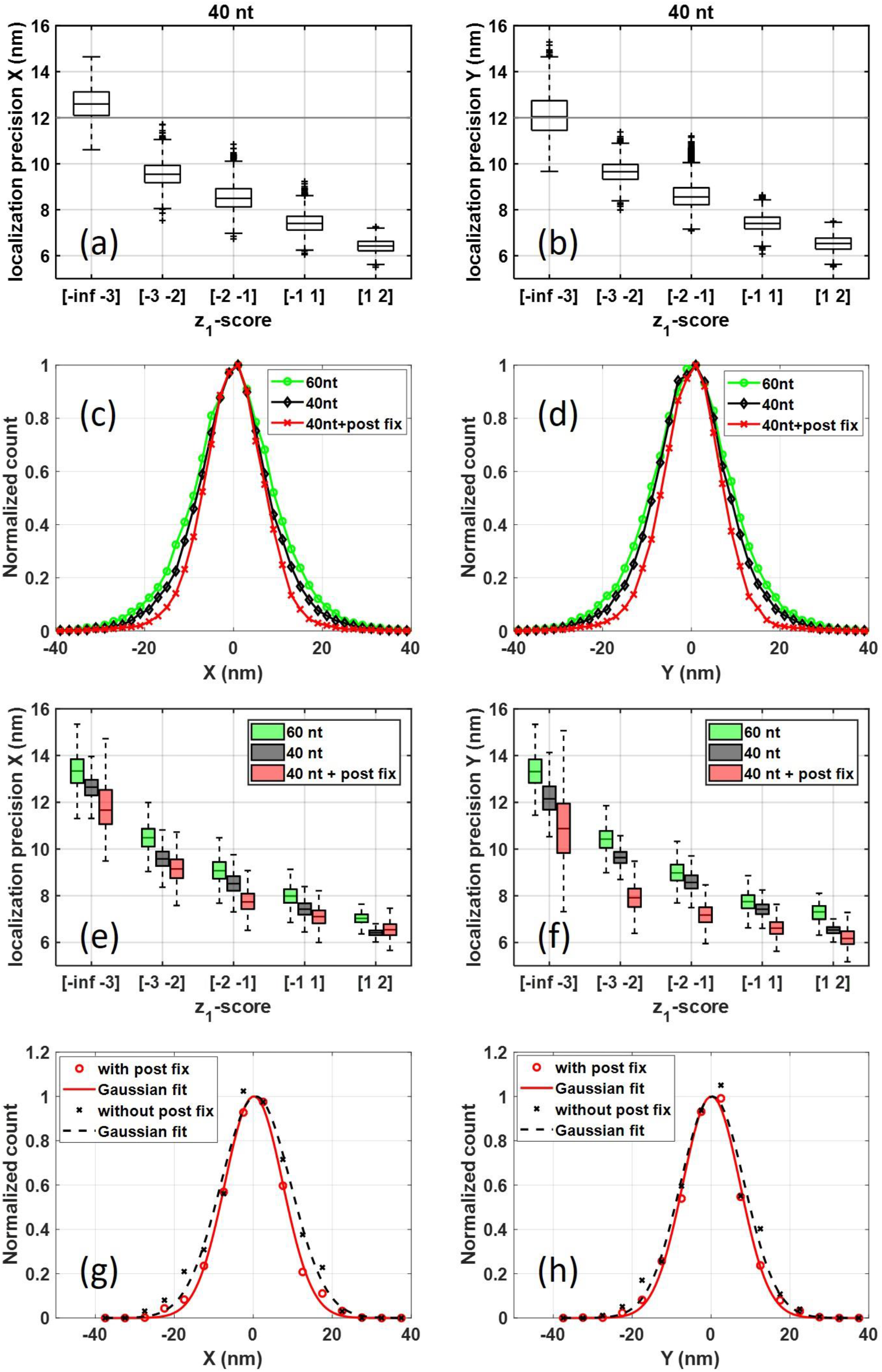
Localization precision in x-direction (left panel) and y-direction (right panel) are affected by the stochastic motion of bio-molecules. (a-b) Boxplot of localization precision distribution calculated from molecules with different *z*_1_-scores. Within each box, horizontal black lines denote median values; boxes extend from the 25^th^ to the 75^th^ percentile of each group’s distribution of values; vertical extending lines denote adjacent values (i.e., the most extreme values within 1.5 interquartile range of the 25^th^ and 75^th^ percentile of each group); (c-d) Comparison of the normalized histogram of localizations from 40-nt oligonucleotides (black line with diamond markers), 60-nt oligonucleotides (green line with circle markers) and post-fixed 40-nt oligonucleotides (red line with cross markers); (e-f) comparison of localization precision of similar *z*_1_-scores from 40-nt oligonucleotides (black), 60-nt oligonucleotides (green) and post-fixed 40-nt oligonucleotides (red); (g-h) Comparison of normalized histogram of localizations from 40-nt oligonucleotides with (red circle) and without (black cross) post fix. The red line shows the Gaussian fit of the experimental data (red circle) from 40-nt oligonucleotides with post fix. The black line shows the Gaussian fit of the experimental data (black cross) from 40-nt oligonucleotides without post fix.

Increasing biomolecule size would increase the range of stochastic motion, leading to further decrease of localization precision. To confirm this, we enlarge the oligonucleotide molecule from 40 nt to 60 nt. The histogram of localizations in x-and y-direction from 2119 molecules with 60-nt size (green line with circle markers) were showed in **Figure 4c** and **4d,** respectively, in comparison of those from molecules with 40-nt size (black line with diamond markers). The histogram data were normalized for better comparison. We can see that, the histogram curves of 60-nt molecules distribute wider than those of 40-nt molecules, suggesting better localization precision.

Adding a ‘post-fix’ step in sample preparation can minimize the bio-molecule motion. To test this, the localization precision of 8840 molecules with 40-nt size with paraformaldehyde cross-linking post fixation (red line with cross makers) were also calculated in **Figure 4c** and **4d**, both of which show the distribution narrower than the curves without post fix.

When comparing molecules of similar z-scores from different samples, we found that in regard to each certain z-score range the localization precisions using 40-nt molecules (black) are all better than those using 60-nt molecules (green, **Figures 4e – 4f**) but not as good as those using samples with post fixation (red, **Figures 4e – 4f**). Therefore, these results reveal the impact of biomolecule stochastic motion on increasing localization uncertainty, providing potential solution for high resolution via reducing probe sizes or further constraining probes.

To further demonstrate improvement of localization precision by minimizing stochastic motion on biology samples, we performed immunofluorescence imaging of microtubules. Two microtubule samples were incubated with mouse anti-tubulin antibody (sigma) and then with Alexa-647-conjugated anti-mouse secondary antibody, with labelling density low enough to ensure that localizations from individual molecules can be identified. One of the samples was post-fixed with 4% paraformaldehyde. The histogram of all the localizations from single molecules from the two samples were plotted in **Figure 4g** (x-direction) and **4h** (y-direction). We can see that the treatment of post fix (red solid line with circle markers) obviously narrowed down the distribution, in comparison of that without post fix (black dashed line with cross markers).

## 3 Conclusion

In summary, we have collected STORM data for immobilized single oligonucleotides and applied *z*_1_-score to evaluate molecule stochastic motion. The *z*_1_-score is supposed to be around zero for immobile molecules as displacements of localizations from the same molecule do not significantly vary along acquisition process. It is expected to be negative for molecules undergoing various motions. In our results, despite of drift correction the main population of oligonucleotides examined were found to exhibit *z*_1_-score of around −1.0, while sample drift was found to contribute to *z*_1_-score with around −1.7 (**Figure 2b and 2c**). These results suggested that confined stochastic motion of immobilized biomolecule is remarkable and detectable in STORM although relatively mild compared to sample drift. Due to dense labeling and stochastic activation, molecules in regular biological samples are untraceable for their complete trajectory in SMLM, possibly accounting for little evidence of confined stochastic motion of immobilized molecules in SMLM. These lines of findings elucidate biomolecule stochastic motion in SMLM. Similar phenomena and impacts are expected in PALM as both techniques provide unmatched super-resolution for biological samples via sequentially resolving single molecules and requiring a relatively long acquisition time [6].

Furthermore, we reveal that molecule stochastic motion facilitates localization uncertainty and decreases localization precision, which is reinforced with increasing molecule sizes. These findings are based on data from immobilized single oligonucleotides containing 40 nucleotides that are of around 12 nm in sizes. In regular practice, antibodies, nucleotide probes or fluorescent proteins are usually used for SMLM. In molecule sizes, these molecules are probably similar to or even larger than the oligonucleotides used in the present study although they might have different robustness. For example, antibodies are around 12 nm, and primary antibody followed by secondary antibody in many biological samples likely doubles the complex sizes and facilitates localization uncertainty. Stochastic motion of these molecules in imaging buffer is expected to increase localization uncertainty in STORM. Due to the different robustness of various biomolecules and different binding/conjugation, for example, primary antibody attached to cellular episode via xxx, more detailed assessments are needed to be carried out in future.

Moreover, this study implies two potential experimental approaches, narrowing down molecule sizes and adding a post-fixation step in sample preparation, to enhance localization precision. Other potential strategies to reduce stochastic motion, such as cooling down samples or adding special ionize into the imaging buffer, could be tested in future experiment to achieve better SMLM resolution. Therefore, this study demonstrates the existence and impact of molecule stochastic motion in SMLM and provides new direction for enhancing localization precision in SMLM.

## 4 Materials and methods

### 4.1 System

A STORM system based on an inverted optical microscope (IX-71, Olympus) with a 100x oil immersion objective lens (Olympus) was used for the nano-imaging as previously described [20]. Astigmatism imaging method was adopted for 3D-STORM, in which a weak cylindrical lens (1 m focal length) was introduced into the imaging path. A 641 nm laser (CUBE 640–100C;Coherent) was used to excite fluorescence and switch Alexa-647 to the dark state. The illumination used the highly inclined and laminated optical sheet (HILO) configuration [21]. The laser power densities were approximately 1.45 kW/cm2 for the 641 nm laser unless otherwise indicated. A dichroic mirror (ZT647rdc, Chroma) was used to separate the fluorescence from the laser and a band-pass filter (FF01-676/37, Semrock) on the imaging path was used to filter the fluorescence. Raw images of the fluorescent signals in each nuclear field were acquired with an EMCCD (DU-897U-CV0, Andor) at 33 Hz for 8000 frames. To avoid focal drift, an anti-drift system was used to sustain the focal position within 10 nm during image processing [22].

### 4.2 Buffer

A standard STORM imaging buffer containing 50Mm Tris (pH 8.0), 10mM NaCl,1% b-mercaptoethanol (v/v), 10% glucose (w/v), 0.5 mg/mL glucose oxidase (G2133, Sigma), and 40mg/mL catalase (C30, Sigma) (23-24) was used.

### 4.3 Label Protocal

To study the stochastic motion of 40-nt oligonucleotides, each coverslip was cleaned by sonication for 15-25 min in 100% Ethanol and washed with Mili-Q water. The cleaned coverslip was incubated with gelatin at room temperature for 10min. 1μM Alexa-647-conjugated oligonucleotides (each 40 nt oligonucleotide labeled with one Alexa-647 at its 3’ end and one biotin at its 5’ end) were adsorbed to the coverslip for 30 min and then incubated with 0.2 μm red fluorescent microspheres (F8810, Thermo Fisher) for 9 min as fiducial markers to correct for sample drift in the x-y plane during imaging. The sample were rinsed extensively with PBS to remove unbound probes before imaging.

To compare the localization precision for different molecular size and post-fixed samples, fiducial markers were incubated before Alexa-647-conjugated oligonucleotides(40 nt or 60 nt). Only post-fixed samples were treated with 4% paraformaldehyde in PBS and washed with PBS 3times and then all the samples were imaged in turn.

To further study the localization precision of cellular microtubules with and without post fix, the immunostaining procedure for microtubules consisted of fixation for 15 min at room temperature containing 3.7% formaldehyde, 0.3% Triton X-100, 0.1% Glutaraldehyde,80mM PIPES pH 6.8, 1 mM EGTA, and 1 mM MgCl2.The fixed cells were rinsed with PBS 3 times, reducted for 7 min with fresh prepare 1mg/ml sodium borohydride to reduce background fluorescence, andwashed with ddH2O 3 times.B The cells were blocked with 2% BSA-PBS for 0.5-1 hour andincubated with mouse anti-tubulin antibody (sigma) in 2% BSA-PBS for 1~2h at room temperature, washed with PBS 3 times (5min per time), and then incubaed with Alexa-647-conjugated anti-mouse secondary antibody at various concentrations for 1 hour at room temperature, washed with PBS at room temperature 3 times.Samples with fixation after labeling were treated with 4% paraformaldehyde in PBS and washed with PBS 3 times (contrasted samples without this step). Finally, samples were embedded in STORM imaging buffer for analysis.

### 4.4 Image processing

For imaging data analysis, a freely available plugin for ImageJ named Thunderstorm was applied to analyze raw images. The precise localization data were from point-like objects in the samples, which appeared as small clusters of localizations. Each cluster contained more than 8 localizations. The clusters in cellular samples were away from any discernable microtubule filaments.

### 4.5 Simulation of confined stochastic motion

Simulation of a 2D stochastic motion of 200 particles within a domain restricted by a radius *R* was performed. The radius is set as 14.6 nm, which is the sum of the length of 40-nt oligonucleotide (40 *nt* × 0.34 *nm/nt*) and the radius of the dye molecule (approx. 1 nm). The position was measured with a constant time interval Δ*t* = 30 *ms* for 8000 frames. The initial position is supposed to be (0, 0). The increment of position for each step is independent and follows a normal distribution, as described by

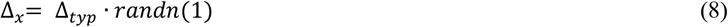

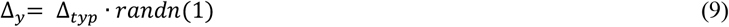

Here, Δ_*typ*_ is the typical movement distance of a molecule during the time interval Δ*t*. The *randn*(1) generates a random number from the standard normal distribution.

The position recorded at each frame can be calculated from their increments,

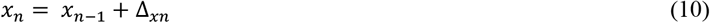

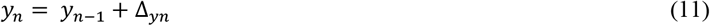

Positions outside the circular domain will be discarded and regenerated by Eq. (8-11).

We assume that the only effect the oligonucleotide has on the dye molecule is to exert a restoring force that increases linearly with the distance of the bead from the attachment point and is directed towards the attachment point. We model the force as a simple spring force 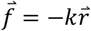. The spring force moves the dye molecule towards to the center with a speed 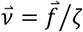, where *ζ* is the dye molecule’s friction constant in imaging buffer. Then we can get the position of the dye molecule by superimposing an additional term 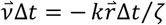 on the random steps of Eqs. (10-11),

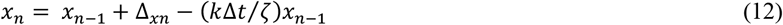

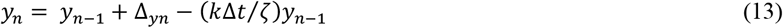

To further considering the localization error, we assume a static localization precision *δ_static_.* The localizations were calculated from Eqs. (12-13) as,

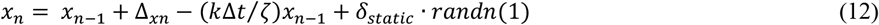

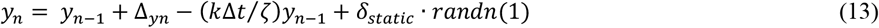

To simulate the sparse activation of AF647, we extracted the on/off pattern of frame sequence of 2073 molecules from our experiment. In our simulation of 2000 molecules, the on/off pattern of each molecule was randomly selected from the experimental data. Eqs. (12-13) were revised to

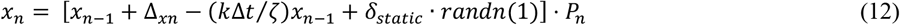

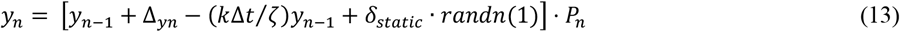

with *P_n_* = 1 standing for on-state and *P_n_* = 0 for off-state.

Eqs. (12-13) generate localizations for an individual molecule undergoing confined stochastic motion. Its *z*_1_-score and *z_m_*-score can be calculated as described in **Section 2.2**.

Three unknown parameters (Δ_typ_, *k*′ = *k*Δ*t/ζ, δ_static_*) were chosen to ensure the output of simulation agree with the experimental data, generating best fit values of Δ_*typ*_ = 5.1769 *nm, k*′ = 0.1257 and *δ_static_* = 5.5 *nm.*

The simulated localization precision from 2000 molecules are 9.6348 nm in x-direction and 9.5143 nm in y-direction, in good agreement with our experimental localization precision of 2073 molecules (40-nt oligonucleotides with biotin-binding), which are 9.0365 nm in x-direction and 9.1096 nm in y-direction.

The simulated curve of *z_m_*-score with different frame is plotted in **Figure 3b**. To reduce running time, 200 molecules are simulated in this part. The saturation behavior is in close analogy to that of experiment in **Figure 3a**. The simulated curve starts with *z*_1_-score of −1.1 equal to that from experiment.

The outputs of localization precision, *z_m_*-score and *z*_1_-score match well with experimental data, suggesting the model work well.

## Acknowledgments

This work was supported by Guangdong Major Project of Basic and Applied Basic Research (No. 2020B0301030009), by National Natural Science Foundation of China (31401146), Shenzhen Science and Technology Planning Project (No. JCYJ20170817095211560), Shenzhen Peacock Plan KQTD20170330110444030; Natural Science Foundation of Guangdong Province: 2016A030312010. X. Yuan appreciates the support given by the leading talents of Guangdong province: No. 00201505.

## Author contributions

The manuscript was written through contributions of all authors. All authors have given approval to the final version of the manuscript.

## Notes

### Competing Interest Statement

The authors have declared no competing interest.

